# Discovery of a novel chemotype targeting *Mycobacterium tuberculosis* cytochrome *bd* through rapid screening and structural elucidation

**DOI:** 10.64898/2026.05.21.726858

**Authors:** Tijn T. van der Velden, Asad Halimi, Julia P. V. Pols, Wing-Sam Lam, Stephan M. Hacker, Lars J. C. Jeuken

## Abstract

Antibiotic resistance in *Mycobacterium tuberculosis* is a pressing global health challenge demanding new therapeutic strategies. The bacterial respiratory chain comprises promising antibacterial targets, with dual inhibition of the terminal oxidases cytochrome *bcc:aa*_*3*_ and cytochrome *bd* (cyt *bd*) showing bactericidal activity. While *bcc:aa*_*3*_ inhibitors such as **Q203** have advanced clinically, cyt *bd* remains underexplored due to difficulties in assigning activity of the purified enzyme and structurally resolving the quinol substrate binding site. Here, we report a rapid *in vitro* screening platform for cyt *bd* inhibitors by engineering a minimal respiratory system that couples the activity of cyt *bd* to that of a type 2 NADH dehydrogenase. This coupled assay enables spectroscopic monitoring of NADH oxidation as a proxy for cyt *bd* activity, allowing rapid screening of over 10,000 compounds. Screening identified **WSL017**, a fragment with low micromolar potency against both *M. tuberculosis* and *E. coli* cyt *bd*. Kinetic and structural analyses revealed competitive inhibition at the quinol-binding site, providing the first structural insights into cyt *bd* inhibition by a non-quinone scaffold. **WSL017** displayed growth inhibition of *M. tuberculosis* H37ra, corroborating oxidase inhibition as a promising therapeutic strategy. This work establishes a pipeline for cyt *bd* inhibitor discovery and highlights new opportunities for structure-guided drug development against cytochrome *bd* oxidases.

## Introduction

Antibiotic resistance is an escalating global health crisis, projected to cause 8.2 million deaths annually by 2050 ^1^. Among resistant pathogens, *Mycobacterium tuberculosis* poses a particularly urgent challenge as it gains antibiotic resistance at the highest pace of all major pathogens ^1^. Tuberculosis (TB) treatment currently requires a prolonged multi-drug regimen lasting up to nine months, imposing significant burden on cost, toxicity, and patient compliance ^2^. A major breakthrough in TB treatment was marked in 2012 with the approval of bedaquiline, shortening treatment duration to six months by targeting the ATP synthase ^3^. However, resistance to bedaquiline has already emerged, driven by an efflux mechanism that expels the drug from the bacterial cell ^4^. These developments underscore the urgent need for next-generation TB antibiotics in order to decrease treatment duration and maintain potent treatment options.

The bacterial respiratory chain offers a compelling target for drug discovery due to its central role in energy metabolism and pathogen survival ^5^ (**Fig 1**). **Q203**, an inhibitor of the terminal oxidase cytochrome *bcc:aa*_*3*_, has successfully completed phase IIa clinical trials ^6^. Yet its bacteriostatic activity reflects metabolic compensation by the alternative terminal oxidase, cytochrome *bd* (cyt *bd*), which sustains electron flow under stress conditions ^7,8^. This respiratory flexibility highlights the necessity of dual inhibition of cytochrome *bcc:aa*_*3*_ and *bd* to achieve bactericidal effects ^9^. Despite its pivotal role, cyt *bd* remains therapeutically underexplored, with no inhibitors currently in clinical development.

**Figure 1.**
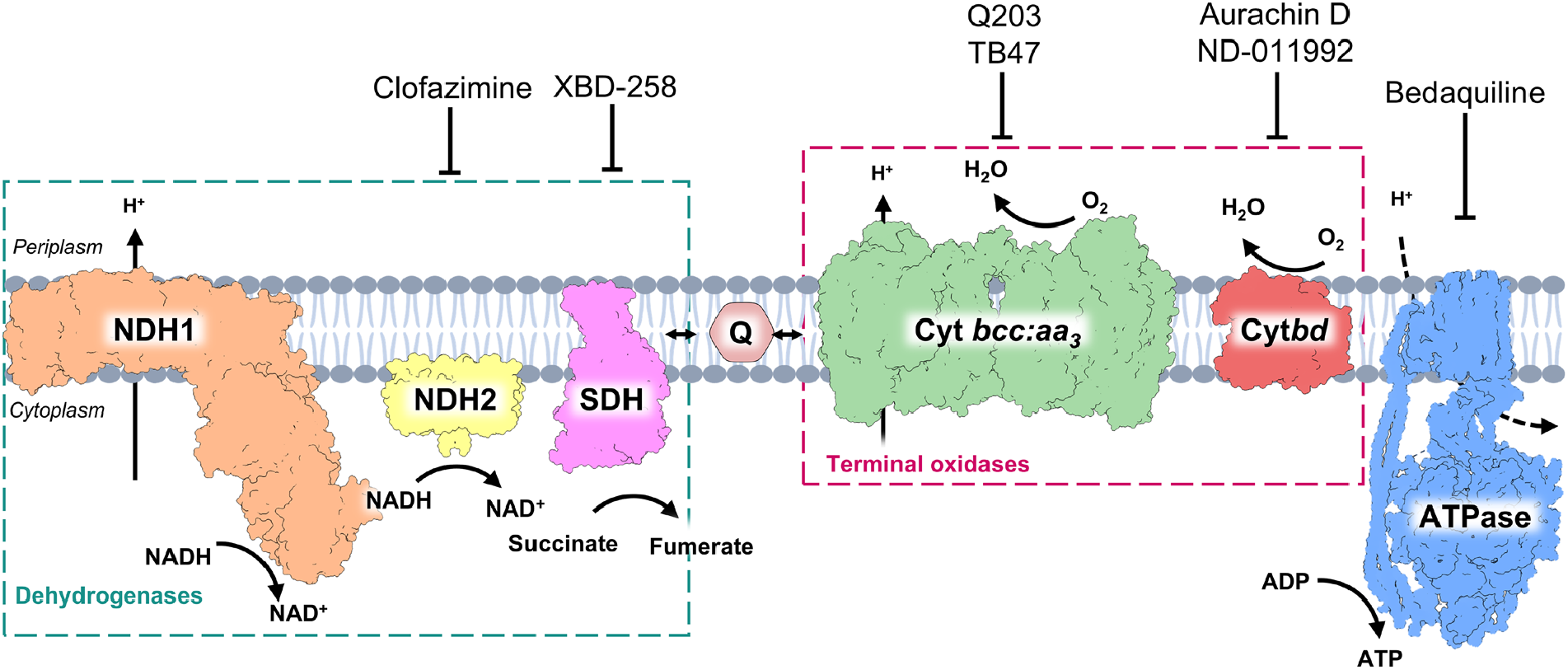
The *M. tuberculosis* respiratory chain and its inhibitors. The dehydrogenases are highlighted in the blue box, and the terminal oxidases in the pink box. Dual inhibition of the two terminal oxidases is highlighted as potent antibiotic strategy. Inhibitors are annotated based on Adolph, C. *et al*.^*31*^ and Saha, P. *et al*.^*32*^. The membrane was generated using Biorender.

Cyt *bd* is distinguished by its high oxygen affinity, enabling respiration under hypoxic conditions typical of infection ^10,11^. Beyond energy generation, cyt *bd* mitigates reactive oxygen species, promotes biofilm formation, and enhances persistence, functions that collectively aid in virulence and antibiotic tolerance ^12–22^. Studies confirm its essentiality during infection and its ability to compensate for inhibition of other respiratory complexes ^8^. Strikingly, cyt *bd* knockout strains exhibit hypersensitivity to bedaquiline, reinforcing its synergistic potential with other respiratory enzymes ^23^. Importantly, cyt *bd* is absent in eukaryotes, making it an attractive target for selective antimicrobial strategies.

Despite this promise, progress on developing cyt *bd* inhibitors has been limited. Existing inhibitors, such as the quinone analog Aurachin D (AurD), exhibit potent inhibition but suffer from cytotoxicity due to structural similarity to ubiquinone, the central respiratory substrate used in mitochondria ^24^. Few synthetic cyt *bd* inhibitors have been reported and often mimic the AurD architecture, potentially maintaining its cytotoxic effects ^25–27^. Furthermore, structural insights remain restricted to AurD bound to *E. coli* cyt *bd (Ecbd)* ^*28,29*^. Expanding the toolbox of cyt *bd* inhibitors and elucidating inhibitor binding modes are critical next steps for structure-guided drug design against the *bd* oxidases.

Thus far, conventional screening approaches have fallen short in populating the drug development pipeline due to technical challenges. *In vivo* assays in *M. tuberculosis* are hampered by slow growth rates and the need for dual oxidase inhibition to achieve bactericidal effects. Additionally, *in silico* approaches are constrained by uncertainty surrounding the *M. tuberculosis* cyt *bd* (*Mtbd)* active site architecture. Substrate and inhibitor bound structures could still not be obtained, even when applying vast molar excess of these ligands, affecting guided docking efforts ^30^. Lastly, current *in vitro* methods studying purified *Mtbd* activity measure its oxygen consumption rate on an Oxygraph system and lack the throughput for large-scale screening.

To overcome these barriers, we developed a rapid *in vitro* screening method for cyt *bd* inhibitor discovery. By engineering a minimal respiratory system incorporating *Caldalkalibacillus thermarum type 2 NADH dehydrogenase* (NDH2) and *Mtbd* (**Fig 2A**), we enable spectroscopic monitoring of NADH oxidation as a proxy for *Mtbd* activity. Using this assay, a library of fragments harboring covalently reactive groups (warheads) was screened. This platform facilitated screening of over 10.000 compounds and yielded a potent inhibitor fragment, which was validated to bind in a non-covalent fashion. Subsequent kinetic and structural characterization of this fragment revealed new opportunities for targeting *bd* oxidases and advancing next-generation TB therapeutics.

**Figure 2.**
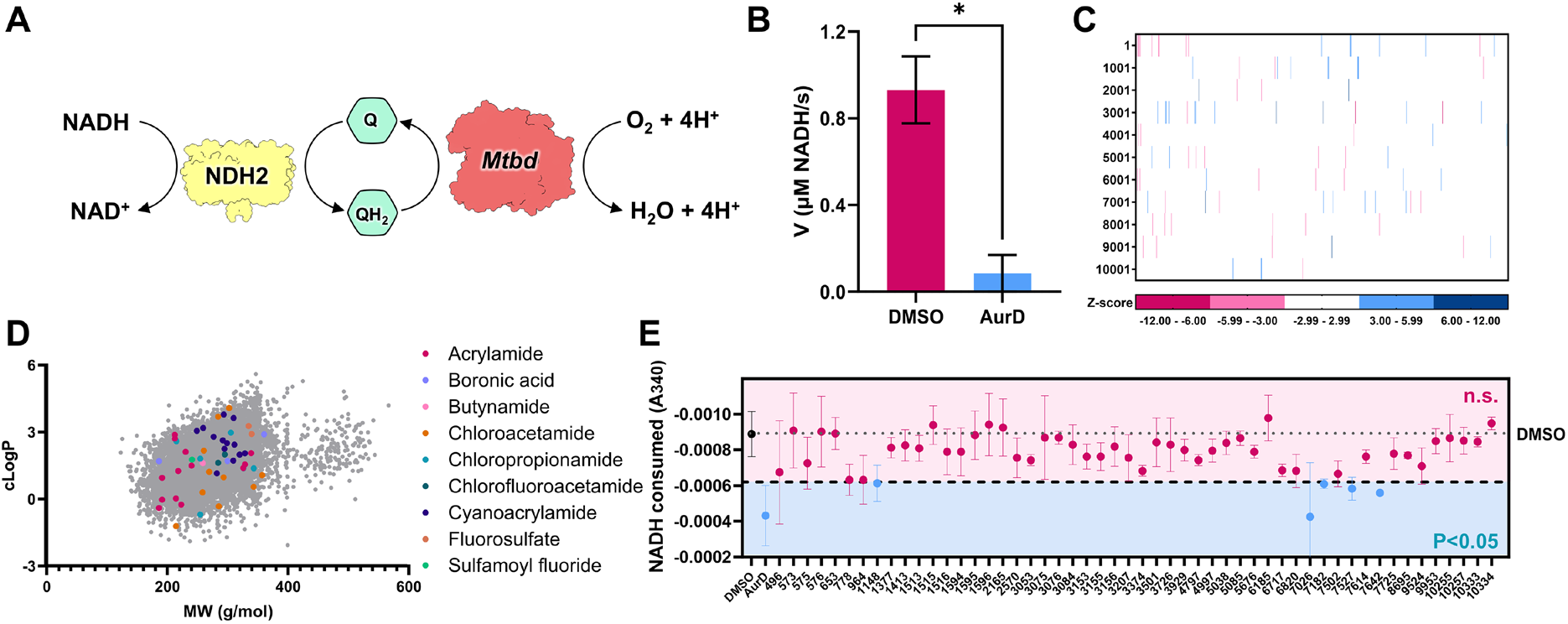
Rapid inhibitor screening strategy against *Mtbd*. (**A**) Overview of the minimal respiratory chain used for inhibitor screening, enabling monitoring of NADH oxidation as a proxy of *Mtbd* activity. (**B**) NADH oxidation rate of the coupled enzyme system showing the inhibitory effect of AurD (1 nM *Mtbd*, 60 nM NDH2, 50 µM menaquinone-1 (MK-1), 250 µM NADH in 50 mM MES pH 6.5, 150 mM NaCl, 0.005% LMNG). (**C**) Heatmap of the inhibitory and stimulatory activity of the full library screened against *Mtbd* at a concentration of 3 µM. (**D**) Hit distribution across the chemical library. Hit compounds are colored by their chemotype. All non-hits are shown in grey. (**E**) Rescreening of the inhibitor hits in *Mtbd* proteoliposomes (POPE:POPG:Cardiolipin; 30:60:10 molar ratio), probing effective diffusion of the inhibitor into the membrane to target *Mtbd* (6 nM *Mtbd*, 60 nM NDH2, 50 µM MK-1, 250 µM NADH, 50 mM MES pH 6.5, 150 mM NaCl, 3 µM Inhibitor). All compound numbers are those of the combined ENL library.

## Results

### Development of a rapid screening method against *Mtbd*

To enable *in vitro* examination of *Mtbd*, the enzyme was heterologously expressed in *M. smegmatis* and purified *via* 3×FLAG affinity chromatography followed by gel filtration (**Fig S1**). Probing of *Mtbd* oxygen consumption was achieved by our previously reported minimal respiratory chain system using excess *C. thermarum* NDH2 as a quinone reductase, enabling the monitoring of *Mtbd* activity without affecting its regulatory disulfide bond ^*33*^ (**Fig 2A**). To improve experimental throughput of this system, we hypothesized that the NADH oxidation rate is representative of *Mtbd* activity, provided *Mtbd* activity is rate limiting, allowing for spectroscopic activity measurements in plate-based assays.

As proof of concept of this system, molar excess of NDH2 was used to maintain the quinone pool reduced and ensure rate limitation by *Mtbd*. The NADH oxidation reaction rate of our minimal respiratory chain was almost completely inhibited upon addition of the known *Mtbd* inhibitor Aurachin D (AurD) (**Fig 2B**), validating this approach for inhibitor screening. Importantly, this spectroscopic assay enables high-throughput plate-based monitoring, in contrast to Oxygraph measurements that quantify *Mtbd* oxygen consumption ^33^.

To discover new chemical scaffolds for cyt *bd* inhibitors, we screened a library of fragments with covalent warheads comprising 10,570 compounds that span diverse sizes and have hydrophobicity properties suitable for membrane-embedded targets. The library consists solely of fragments with covalent warheads in order to increase the chance of finding initial hits. Screening in the coupled assay at 3 µM compound concentration yielded 52 primary inhibitor hits (0.49% hit rate) (**Fig 2C, Table S1**), and 43 stimulator hits. For the inhibitors, 48 hits were confirmed upon triplicate testing, while none of the stimulators showed significant effects.

Inhibitor hits were broadly distributed across size, cLogP, and chemotype (**Fig 2D**). To ensure activity in a native-like membrane environment, hits were re-screened in proteoliposomes reconstituted with our minimal respiratory chain system. Five compounds retained significant inhibitory activity at 3 µM concentration (**Fig 2E**). Among these, a chloroacetamide derivative, compound **ENL1148** (Z57044282), exhibited a clear dose-response relationship and was prioritized for further characterization.

### Inhibitor ENL1148 is competitive and acts through a non-covalent mechanism

Despite its covalent warhead, washout and time-based inhibition assays indicated that **ENL1148** does not inhibit *Mtbd* covalently. To confirm this, we synthesized a non-covalent analog lacking the chloroacetamide warhead, yielding compound **WSL017** (**Fig 3A, Fig S2,S3**). Further kinetic evaluation of the inhibitors was performed measuring the cyt *bd* oxygen consumption rate on an Oxygraph system. Both **ENL1148** and **WSL017** displayed low micromolar IC_50_ values against *Mtbd* and *Ecbd* in our proteoliposomal oxygen consumption assays (**Fig 3B,C**), demonstrating potent activity for a small fragment. To ensure that the observed inhibition was acting directly on the *bd* oxidase, the activity of NDH2 was measured in a separate assay. Neither **ENL1148** nor **WSL017** showed inhibition of NDH2 activity (**Fig 3D**), confirming the direct inhibition of *Ecbd* and *Mtbd*. Further experiments were performed using the non-covalent compound **WSL017**. Kinetic analysis of **WSL017** revealed competitive inhibition of *Mtbd* and *Ecbd*, significantly increasing *K*_*M*_ without a significant effect on V_max_ (**Fig 3E,F**). These data suggest that **WSL017** binds at the quinol-binding site, displacing the natural quinol substrate in both *Mtbd* and *Ecbd*.

**Figure 3.**
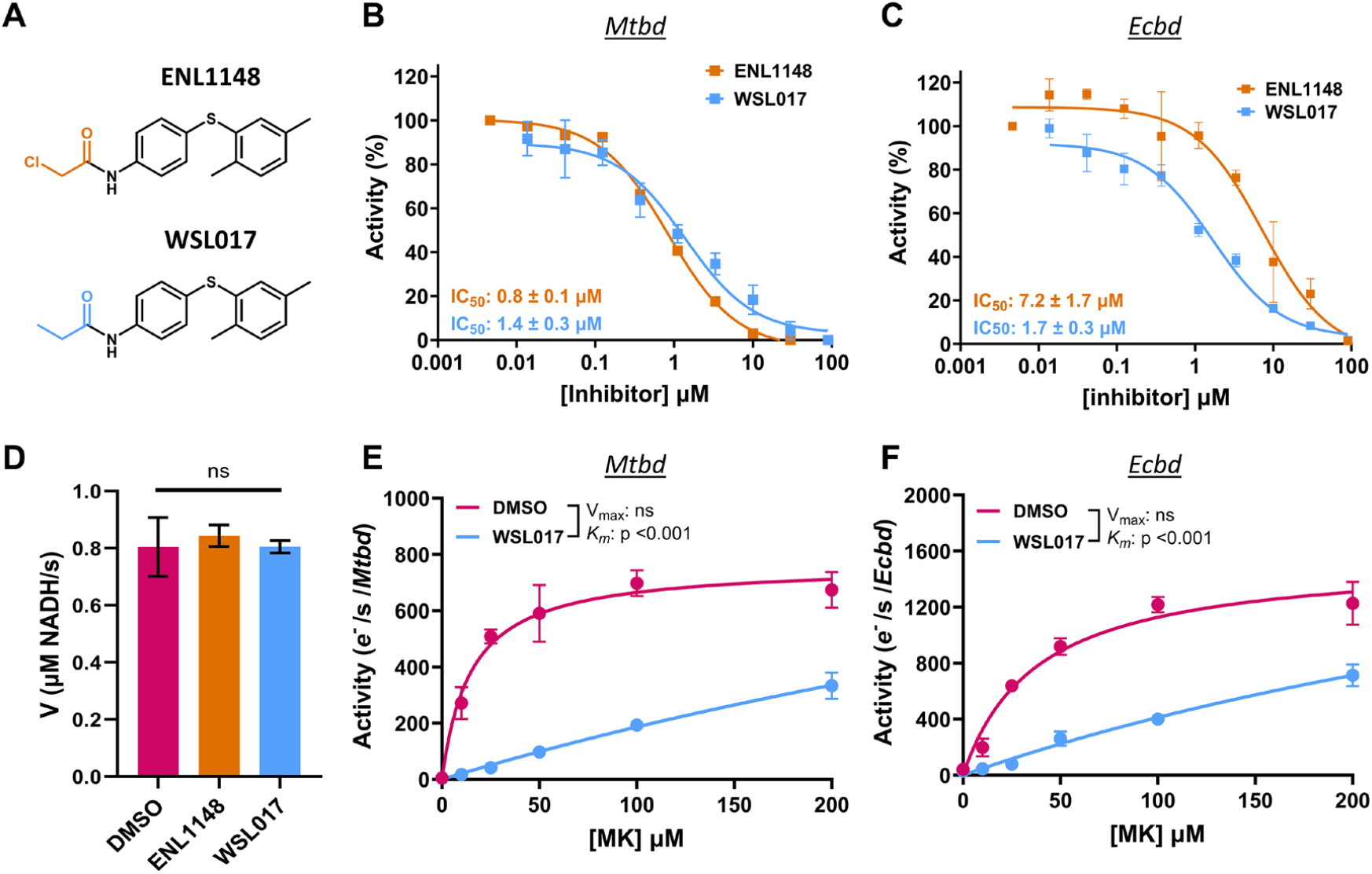
Inhibitor kinetics of ENL1148 and WSL017. (**A**) Inhibitor structure of **ENL1148** and **WSL017**, highlighting the removal of the warhead. (**B**) IC_50_ determination on the Oxygraph system against *Mtbd* proteoliposomes (POPE:POPG:Cardiolipin ; 30:60:10 molar ratio) and (**C**) against *Ecbd* proteoliposomes (POPE:POPG:Cardiolipin ; 60:30:10 molar ratio). (**D**) Neither inhibitor affects NDH2 activity (100 nM NDH2, 200 µM menadione, 150 µM NADH). (**E**) Inhibitor competition kinetics against *Mtbd* proteoliposomes and (**F**) *Ecbd* proteoliposomes. Assays were performed using 6 nM cyt *bd*, 60 nM NDH2, 50 µM MK-1, 50 mM MES pH 6.5, 150 mM NaCl.

### Cryo-EM reveals the binding mode of WSL017 in the *Ecbd* active site

To confirm the binding site of **WSL017**, we conducted cryogenic electron microscopy (cryo-EM) on *Mtbd. Mtbd* was plunge frozen in the presence of **WSL017**, and under turnover conditions in the presence of **WSL017** and was solved between 2.9 Å and 3.2 Å resolution. The overall *Mtbd* structure showed to be identical to the one reported before ^30^, consisting of the subunits CydA and CydB. Regretfully, none of these conditions showed a density that could be assigned to **WSL017**. Neither did the resulting density maps show displacement of the menaquinone (MK) bound at Trp9^CydA^ (**Fig S4**). Similar observations have been made before, even after incubation with vast molar excesses of the canonical inhibitor AurD ^30^. The lack of ligand binding, whether substrate or inhibitor, to *Mtbd* hinders drug development and remains subject to further study.

To still gain insights into the binding mode of **WSL017** to *bd* oxidases, we performed cryo-EM of *Ecbd* in the presence of **WSL017**. The structure of *Ecbd* was solved in both a monomeric and dimeric state at 2.64 Å and 2.58 Å, respectively (**Fig S5-S7**). The *Ecbd* structures resemble previously reported structures, consisting of the subunits CydA (including the quinol binding Q-loop), CydB, CydX, and CydH, and the characteristic triangular heme arrangement containing heme *b*_*558*_, *b*_*595*_ and *d* ^*29,34*^. As seen before, the flexible Q-loop lid (Glu257^CydA^ to Gly306^CydA^) in the monomer state of *Ecbd* prohibits assignment of any ligand in the quinol oxidation site ^29^. This flexible quinol oxidation site in *bd* oxidases prohibited structure guided drug discovery until recent structural advances showed a dimeric state of *Ecbd* ^*28,29*^.

In the dimeric state of *Ecbd*, the Q-loop is stabilized and allows for modelling of the active site architecture. We previously reported binding of menaquinone and AurD in this dimeric *Ecbd* state, revealing Y243^CydA^ and R298^CydA^ as the main catalytic residues ^29^. Under turnover conditions, the Q-loop lid (Glu257^CydA^-Gly306^CydA^) undergoes a disorder-to-order transition to enable quinol oxidation. Here, **WSL017** was shown to occupy the same quinol oxidation site as menaquinone in the substrate bound state (**Fig 4A**). **WSL017** is bound in a shallow pocket under the Q-loop lid typically unoccupied without added ligands. In this pocket, binding of **WSL017** is stabilized by the sulfur moiety interacting with Y243^CydA^ (**Fig 4B**), that typically forms interactions by π-π stacking with the menaquinone headgroup during substrate binding (**Fig 4C**). This pocket is characterized by hydrophobic interactions with CydA Val236, Leu237, Gly238, Glu240, Tyr243, Phe269, Ile295, Ala296, Arg298, Phe390, and Heme *b*_*558*_. These hydrophobic interactions maintain the Q-loop in the closed state, impairing the disorder-to-order transitions typically seen during quinol turnover ^29^. We have previously shown that the mutation of CydA^Y243A^ abolished enzyme activity ^29^, prohibiting the interrogation of the effect of this mutation on inhibitor resistance. Interestingly, the amide side chain of **WSL017** makes few interactions and points out of the pocket, explaining why the removal of the covalent warhead of inhibitor **ENL1148** had no effect on inhibition.

**Figure 4.**
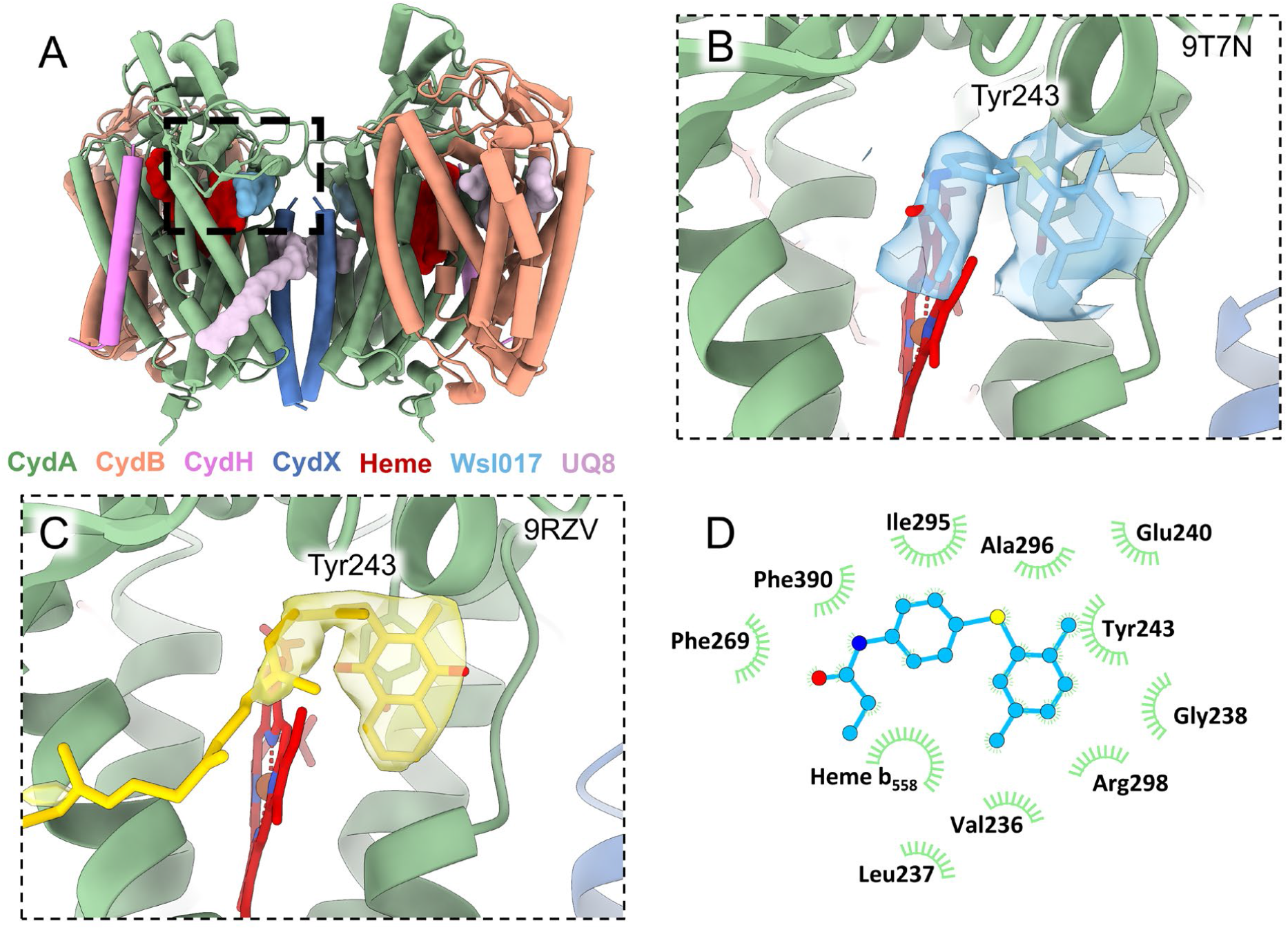
Structural features of *Ecbd* bound to WSL017. (**A**) Cryo-EM structure of the *Ecbd* dimer bound to **WSL017**. (**B**) Density of the active site pocket highlighting **WSL017** and its density feature interacting with Tyr243^CydA^. (**C**) Density of the previously solved active site pocket highlighting a bound menaquinone interacting with Tyr243^CydA^ (PDB 9RZV) ^29^ (**D**) Schematic representation of the **WSL017** binding pocket within subunit CydA quinol oxidation site as analyzed by Ligplot^35^. Hydrophobic interactions are indicated by spoked semi circles.

### WSL017 inhibits growth of *M. tuberculosis* H37ra

To validate the inhibitor strategy with **WSL017** in cells, we performed growth inhibition assays on the attenuated *M. tuberculosis* H37ra, representative of the virulent *M. tuberculosis* H37rv ^36^. To determine inhibition, H37ra was exposed to **WSL017** and/or **Q203** and incubated at 37ºC for 7 days. Due to the low cell densities, growth was determined using Alamar blue, allowing for fluorescent capture of the metabolic activity of remaining cells ^37^.

Upon exposure to 25 µM **WSL017**, H37ra showed significant growth impairment, although not achieving complete growth inhibition (**Fig 5**). Exposure to the Cyt *bcc:aa*_*3*_ inhibitor **Q203** at 17 nM showed no significant effects, while supplementation of **WSL017** with 17 nM **Q203** showed marginally, but not significantly reduced growth compared to **WSL017** alone.

**Figure 5.**
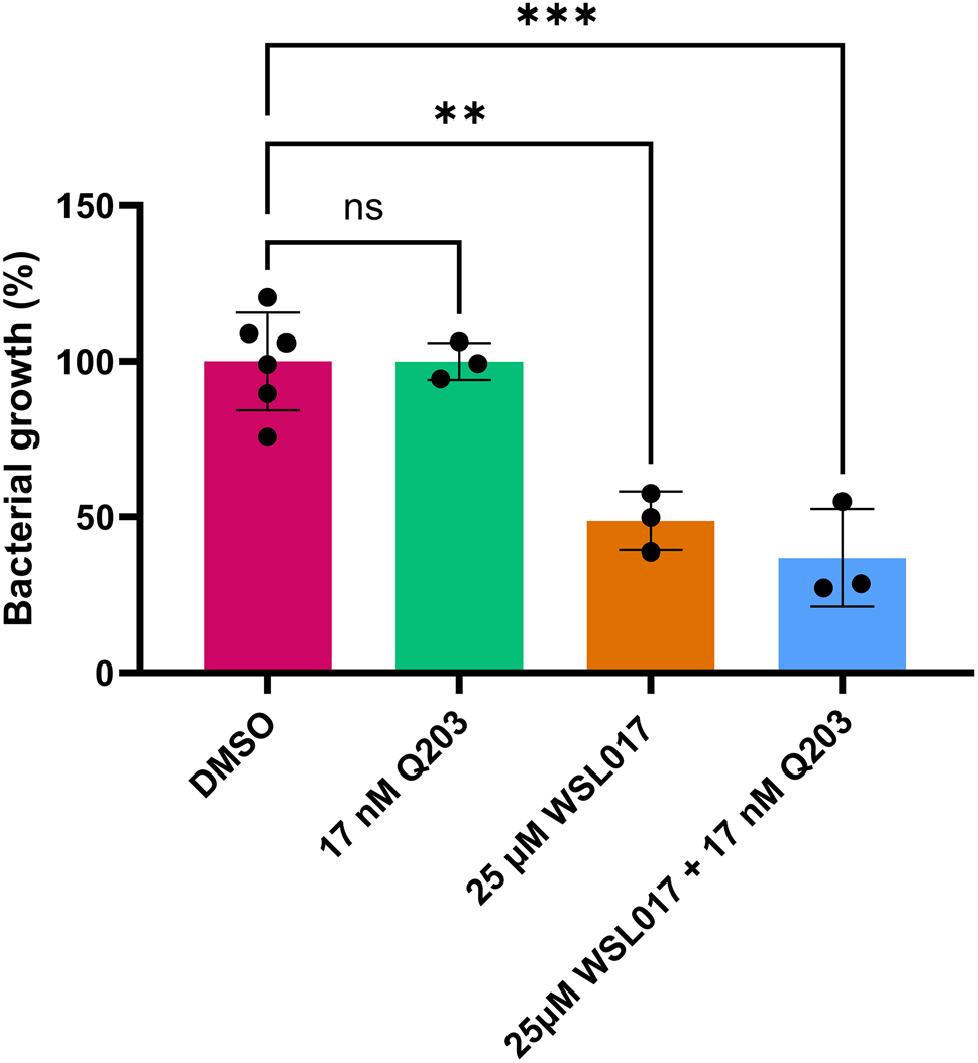
*In cellulo* inhibition of *M. tuberculosis* H37ra growth by the *bd* oxidase inhibitor WSL017 and *bcc:aa*_*3*_ inhibitor Q203.

## Discussion

Tuberculosis is a major health threat, gaining resistance at a rate outpacing other human pathogens ^1^. Especially for these resistant cases, the respiratory chain of *M. tuberculosis* has been shown as key weakness for antibiotic intervention ^38^. The combinatory inhibition of the two terminal oxidases, cytochrome *bcc:aa*_*3*_ and *Mtbd*, holds major potential for treating tubercular infections ^7^. Here, we present a spectroscopic assay platform that allows for rapid kinetic screening for *Mtbd* inhibitors. Using this engineered minimal respiratory chain, we identified the inhibitor **WSL017**, showing low µM potency against both *Mtbd* and *Ecbd*. Notably, this assay can be broadly adapted to target other respiratory enzymes, including NADH dehydrogenases and quinol oxidases from diverse pathogens such as *Staphylococcus aureus* and *Listeria monocytogenes*, expanding its utility beyond the work presented here ^5^.

It is important to note that in the results presented here, *Mtbd* was assayed in liposomes and did not display substrate inhibition kinetics at the MK-1 concentrations tested, which allowed for the determination of **WSL017** binding competitively to the *Mtbd* quinol oxidation site. This is in contrast with our previous report, where *Mtbd* is subject to substrate inhibition at high MK-1 concentrations when assayed in detergent ^33^. We hypothesize that this difference stems from distinct local substrate concentrations at the *Mtbd* active site due to altered diffusion of the quinone into the liposome or detergent micelle.

Mechanistic and structural analyses revealed that **WSL017** acts as a competitive inhibitor, displacing quinol at the cyt *bd* active site. Cryo-EM of *Ecbd* provided the first structural insights into *bd* oxidase inhibition by a non-quinone scaffold, highlighting a hydrophobic pocket that stabilizes **WSL017** and offers opportunities for rational optimization. Optimization might utilize π-π stacking to Phe269^CydA^ and Phe390^CydA^, together with hydrogen bonding to Arg298^CydA^ and the conserved Glu240^CydA^. Especially interactions with Glu240^CydA^, Tyr243^CydA^ and Arg298^CydA^ will prove valuable due to the conservation and essentiality for oxidase activity in *Ecbd* ^29^. Interestingly, the inhibitor binding pocket here is not conserved in *Mtbd*, indicating that **WSL017** binding is mediated by different interactions or occurs at a different pocket (**Fig S8**). Although no ligand density was observed in our *Mtbd* maps, these findings underscore the feasibility of targeting *bd* oxidases with novel chemotypes.

**WSL017** shares structural similarity with previously reported *bd* oxidase inhibitors ND-011992 ^25^ and CK-2-63 ^26^, but lacks the AurD headgroup implicated in cytotoxic side effects (**Fig S9**) ^24^. Surprisingly, the absence of this moiety in **WSL017** indicates that it is not required for binding to the cyt *bd* active site. Unfortunately, no structure has been solved to date with either CK-2-63 or ND-011992 bound to *Ecbd*, or any other *bd* oxidase, preventing structural comparison to **WSL017**. Since **WSL017** is similar to these two inhibitors, but without the AurD type headgroup, it will be interesting to see, how each inhibitor feature contributes interactions in the *Ecbd* binding pocket.

Excluding the AurD quinolone moiety from the next generation of cyt *bd* inhibitors may reduce cross-reactivity with human respiratory enzymes, improving selectivity. Furthermore, unlike AurD, which induces active-site refolding in *Ecbd* CydA helix 6 by hydrogen bonding with Asp239^CydA 29^, **WSL017** inhibits *Ecbd* without triggering such conformational changes, showing a distinct competitive mechanism of action (**Fig S10**). **WSL017** binds *via* predominately hydrophobic interactions within the main body of CydA and the Q-loop lid, maintaining the Q-loop lid in a closed state. We hypothesize that this prohibits the disorder-to-order transitions characteristic of *Ecbd* quinol oxidation, impairing release of **WSL017**. This indicates that the structural changes induced by AurD are not required for inhibition of cyt *bd* and opens a new chemical space for cyt *bd* inhibitor development.

Despite its low µM IC_50_ value *in vitro*, **WSL017** did not yield complete growth inhibition *in cellulo*. In the H37ra strain assayed here, **WSL017** reached growth inhibition of approximately 50%, despite complementation with the cytochrome *bcc:aa*_*3*_ inhibitor **Q203**. This is in stark contrast with inhibitors such as ND-011992 and CK-2-63, which are bactericidal in combination with **Q203** ^7,26^. This indicates that **WSL017** requires further optimization to reach improved affinity or local concentrations in the mycobacterial inner membrane to better inhibit the *bd* oxidase.

Future work should focus on optimizing **WSL017** to enhance potency and studying its specificity, as well as evaluating its activity against *bd* oxidases in other clinically relevant pathogens. Given the demonstrated synergy between *bd* and *bcc:aa*_*3*_ inhibition ^7^, such compounds could form the basis of combination therapies that shorten treatment duration and overcome resistance in tuberculosis and beyond.

### Experimental section

#### Chemistry

##### General remarks

All reagents and solvents were purchased from commercial sources and were of analytical grade (TCI, Sigma-Aldrich and BroadPharm^®^). All moisture sensitive reactions were performed under inert atmosphere. Solvents were dried using 4 Å molecular sieves prior to use, when anhydrous conditions were required. Water used in reactions was always demineralized. Analytical Thin-layer Chromatography (TLC) was routinely performed to monitor the progression of a reaction and was conducted on Merck Silica gel 60 F254 plates. Compounds on the TLC plates were visualized by UV irradiation (λ_254_) and/or spraying with potassium permanganate solution (K_2_CO_3_ (40 g), KMnO_4_ (6 g), and H_2_O (600 mL)) followed by heating as appropriate. Purification by flash column chromatography was performed using Screening Devices B.V. silica gel 60 (40-63 µm, pore diameter of 60 Å).

Solutions were concentrated using a Heidolph laborota W8 4000 efficient rotary evaporator with a Laboport vacuum pump. Analytical purity was determined with Liquid Chromatography-Mass Spectrometry (LC-MS) using a Finnigan LCQ Advantage MAX apparatus with electrospray ionization (ESI), equipped with a Phenomenex Gemini 3 μm NX-C18 110 Å column (50 × 4.6mm), measuring absorbance at 254 nm using a Waters 2998 PDA UV detector and the m/z ratio by using an Acquity Single Quad (Q1) detector. Injection was with the Finnigan Surveyor Autosampler Plus and pumped through the column with the Finnigan Surveyor LC pump plus to be analysed with the Finnigan Surveyor PDA plus detector. Samples were analyzed using eluent gradient 10% → 90% ACN in MiliQ water (+ 0.1% TFA (v/v)).

^1^H and ^13^C Nuclear Magnetic Resonance (NMR) spectra were recorded on a Bruker AV 400 (400/100 MHz) spectrometer at ambient temperature using CDCl_3_ as solvent. Chemical shifts (δ) are referenced in parts per million (ppm) with tetramethylsilane (TMS) or CDCl_3_ resonance as the internal standard peak (δ 0.00 for ^1^H (TMS), δ 77.16 for ^13^C (CDCl_3_)). Multiplicity is reported as s = singlet, d = doublet, dd = doublet of doublet, t = triplet, q = quartet, m = multiplet. Coupling-constants (*J*) are reported in Hertz (Hz).

### *N*-(4-((2,5-dimethylphenyl)thio)phenyl)propionamide (WSL017)

4-((2,5-Dimethylphenyl)thio)aniline (50.0 mg, 218 μmol, 1.0 eq.) and TEA (151.7 μL, 1.1 mmol, 5.0 eq.) were dissolved in THF (1 mL) and the solution was cooled to 0°C. Propionyl chloride (38.1 μL, 436 μmol, 2.0 eq.) was added dropwise, and the reaction mixture was stirred overnight at rt. After TLC analysis indicated completion of the reaction, sat. NH_4_Cl_(aq.)_ (10 mL) was added. The mixture was diluted with EtOAc and the organic and aqueous layers were separated. The organic layer was washed with sat. NH_4_Cl_(aq.)_, brine, dried over MgSO_4_, filtered and concentrated *in vacuo*. The crude product was purified with column flash chromatography (0 → 25% EtOAc in pentane) to yield **WSL017** as a white solid (55.1 mg, 193 μmol, 89%). ^1^H NMR (400 MHz, CDCl_3_) δ 7.48 – 7.44 (m, 2H), 7.39 (s, 1H), 7.22 – 7.16 (m, 2H), 7.10 (d, *J* = 7.7 Hz, 1H), 7.03 (s, 1H), 6.98 (d, *J* = 7.7 Hz, 1H), 2.38 (q, *J* = 7.6 Hz, 2H), 2.31 (s, 3H), 2.23 (s, 3H), 1.23 (t, *J* = 7.6 Hz, 3H). ^13^C NMR (101 MHz, CDCl_3_) δ 172.23, 136.91, 136.41, 136.12, 134.24, 132.56, 131.38, 130.70, 130.49, 128.46, 120.63, 30.85, 20.97, 20.13, 9.78. LC-MS (ESI, 10% → 90% ACN in MiliQ water (+ 0.1% TFA (v/v)): t_R_ = 7.96 min; HRMS calculated for C_17_H_19_NOS + H^+^: 286.12601, found 286.12621. HRMS calculated for C_17_H_19_NOS + Na^+^: 308.10796, found 308.10822.

### *M. tuberculosis* cytochrome *bd (Mtbd)* expression and purification

*Mtbd* was expressed and purified as described before^33^. *M. smegmatis* MC2 155 ΔCydAB, transformed with the plasmid pLHCyd, was grown in LB-hygromycin (50 µg/mL for 72 hours at 250 RPM and 37°C. This culture was then diluted 1:100 and grown for an additional 72 hours at 200 RPM and 37°C. Cells were harvested by centrifugation (6371 rcf, 20 min, 4°C) and resuspended in 5 mL of buffer containing 50 mM Tris-HCl (pH 7.4), 5 mM MgCl_2_, 0.05% Tween-80, and cOmplete™ EDTA-free Protease Inhibitor per gram of wet cells. Cell lysis was performed by two passes through a Stansted pressure cell homogenizer at 270 MPa. Debris was removed by centrifugation (10,000 rcf, 20 min, 4°C), and crude membranes were collected by ultracentrifugation (200,000 rcf, 1 h, 4°C). Membranes were resuspended in 20 mM Tris-HCl (pH 7.4), 0.05% Tween-80, and 10% glycerol to a final protein concentration of 10 mg/mL. Solubilization was achieved by adding 0.5% Lauryl Maltose Neopentyl Glycol (LMNG) and gently mixing for 1 hour at 4°C. Insoluble material was removed by ultracentrifugation (200,000 rcf, 30 min, 4°C). The soluble fraction was incubated overnight at 4°C with Pierce™ Anti-DYKDDDDK Affinity Resin (Thermo Fisher) and purified according to the manufacturer’s protocol. Flow-through was removed by centrifugation (1000 rcf), followed by four washes with 10 column volumes of buffer containing 50 mM Tris-HCl (pH 7.4), 150 mM NaCl, and 0.005% LMNG. *Mtbd* was eluted using buffer composed of 50 mM Tris-HCl (pH 7.4), 150 mM NaCl, 0.02% DDM, and 1 mg/mL 3X-FLAG peptide (Genscript). Final purification was performed by size-exclusion chromatography on a Superdex 200 Increase 10/200 column at 0.5 mL/min using buffer containing 50 mM Tris-HCl (pH 7.4), 150 mM NaCl, and 0.005% LMNG. SEC peak fractions were analyzed by SDS-PAGE, and pure *Mtbd* was pooled, concentrated, and stored at –80°C until use.

### *E. coli* cytochrome *bd* expression and purification

*Ecbd* was expressed and purified as previously described ^33^. *E. coli* MB43 cells, transformed with pET17b-CydABX-linkerstreptag, were cultured overnight in LB medium containing 100 μg/mL ampicillin (250 RPM and 37°C). The culture was diluted to an OD of ∼0.1 and grown to OD ∼0.4 before induction with 0.45 mM IPTG. Cells were harvested at OD 2.0 by centrifugation (6371 rcf, 20 min, 4°C) and resuspended in buffer containing 50 mM MOPS (pH 7.4), 100 mM NaCl, and cOmplete™ EDTA-free Protease Inhibitor (Roche) at a ratio of 5 mL per gram of wet cell pellet. Cell lysis was performed using a Stansted pressure cell homogenizer at 270 MPa. Cell debris was removed by centrifugation (10,000 rcf, 20 min, 4°C), and crude membranes were isolated by ultracentrifugation (200,000 rcf, 1 h, 4°C). Membranes were resuspended in 50 mM MOPS, 100 mM NaCl, pH 7.4, to a final protein concentration of 10 mg/mL before solubilization of *Ecbd* was achieved by incubating with 0.5% LMNG for 1 hour at 4°C with gentle mixing. Insoluble material was removed by ultracentrifugation (200,000 rcf, 30 min, 4°C), and the supernatant was loaded onto a StrepTrap HP column (Cytiva) at a flow rate of 2 mL/min. The column was washed with buffer containing 50 mM sodium phosphate, 300 mM NaCl, 0.005% LMNG, pH 8.0, to remove unbound proteins. *Ecbd* was eluted using buffer composed of 50 mM sodium phosphate, 300 mM NaCl, 2.5 mM desthiobiotin, and 0.005% LMNG, pH 8.0. Peak fractions were pooled, concentrated, and stored at –80°C until further use.

### NDH2 expression and purification

To enable measurement of Cyt *bd* activity, NDH2 was required to reduce the quinone pool. Expression and purification of NDH2 followed previously established methods ^33^. In brief, *E. coli* C41 (DE3) cells, transformed with pET28-NDH2_NtermHis, were cultured overnight in LB medium supplemented with 50 µg/mL kanamycin at 250 RPM and 37°C. The overnight culture was diluted 1:20 and grown to an optical density of ∼0.5 before induction with 0.25 mM IPTG. NDH2 expression proceeded for 4 hours at 37°C, after which cells were harvested by centrifugation (6371 rcf, 20 min, 4°C). Cell pellets were resuspended in 50 mM Tris-HCl, 5 mM MgCl_2_, pH 8.0 (5 mL per gram of cells) and lysed using a Stansted pressure cell homogenizer at 270 MPa. Cell debris was removed by centrifugation (10,000 rcf, 20 min, 4°C), and crude membranes were collected by ultracentrifugation (200,000 rcf, 1 h, 4°C). Membranes were resuspended in buffer containing 50 mM Tris-HCl, 150 mM NaCl, and 20 mM imidazole at a concentration of 10 mg/mL total protein. NDH2 was solubilized by incubating the membrane suspension with 1% DDM for 1 hour at 4°C with gentle mixing. Insoluble material was removed by ultracentrifugation (200,000 rcf, 30 min, 4°C), and the supernatant was loaded onto a HiTrap Nickel NTA column (Cytiva). Unbound proteins were washed with buffer containing 50 mM Tris-HCl pH 8.0, 150 mM NaCl, 20 mM imidazole, and 0.02% DDM. NDH2 was eluted using 30% elution buffer (50 mM Tris-HCl, 150 mM NaCl, 500 mM imidazole, 0.02% DDM). Final purification was performed by size-exclusion chromatography on a Superdex Increase 200 10/300 column (Cytiva) at a flow rate of 0.5 mL/min using buffer containing 50 mM Tris-HCl, 500 mM NaCl, 5% glycerol, and 0.02% DDM. Purified NDH2 fractions were pooled, concentrated, and stored at –80°C until use.

### High throughput inhibitor screening

Inhibitor screens were performed in 384-well format using the following reaction solution: 1 nM *Mtbd*, 60 nM NDH2, 50 µM menaquinone-1 in 50 mM MES pH 6.5, 150 mM NaCl, 0.005% LMNG, and 3 µM of the inhibitor candidate (Enamine libraries CFL-8480 and CYS-3200). A surplus of NDH2 (60 nM) was added to ensure rate limitation by *Mtbd*. The reaction was initiated by the addition of 250 µM NADH. The NADH oxidation rates were monitored at 340 nm in a FLUOstar Omega plate-reader (BMG LABTECH). Each plate contained multiple 1 µM Aurachin D positive controls and DMSO solvent controls to ensure assay validity. The sample activity for each well was determined by the initial slope of NADH oxidation. Each inhibitor was ascribed a Z-score according to the following equation, where *X* is the observed reaction rate, *µ* is the plate mean, and *σ* is the plate standard deviation.

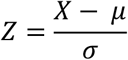

Primary hits were identified with a Z-score < -3.0 (stimulator) or > 3.0 (inhibitor). Hit validation was performed in triplicate, where the inhibitors were compared to the DMSO control. Hits were deemed significant (p<0.05) after ANOVA with Dunetts *post hoc* analysis.

### Cyt *bd* proteoliposomes reconstitution

Cyt *bd* was reconstituted in proteoliposomes based on the method by Gofoy-Hernandez *et al*. ^*39*^. Lipids were obtained from Avanti Polar Lipids and used as received. Lipid mixtures were prepared in CHCl_3_ using POPE:POPG:cardiolipin (CA) in ratios of 60:30:10 for *Ecbd* proteoliposomes and 30:60:10 for *Mtbd* proteoliposomes ^40^. These mixtures were dried under a stream of nitrogen. Residual chloroform was removed overnight by vacuum desiccation. The resulting lipid film was rehydrated to a final concentration of 10 mg/mL lipids in buffer containing 20 mM MOPS, 30 mM Na_2_SO_4_, and 100 mM KCl at pH 7.4, using vortexing to ensure thorough resuspension. Cyt *bd* in LMNG (0.00005% final concentration) was added to the liposome suspension at a protein-to-lipid ratio of 1% (w/w) and mixed by inversion for 30 minutes at room temperature. Insoluble material was removed by centrifugation using an Eppendorf tabletop centrifuge (14,100 rcf, 5 min).

To determine the concentration of cyt *bd* in the proteoliposomes, a sample was solubilized in 2% octyl-β-glucoside and analyzed by UV-visible spectroscopy. Quantification was based on the soret band using the appropriate extinction coefficients: *Ecbd* at 417 nm (ε = 230 mM^−1^ cm^−1^) ^41^ and *Mtbd* at 414 nm (ε = 279 mM^−1^ cm^−1^) ^33^. NDH2 was added to the liposomes at a 10:1 molar ratio NDH2:Cyt *bd* and self-associated to the membrane.

### Hit confirmation and characterization in proteoliposomes

Confirmed hits were further investigated by screening for inhibition in *Mtbd* proteoliposomes, ensuring the inhibitors can diffuse into the membrane to reach the binding site. Proteoliposomes were assayed with 6 nM *Mtbd* or *Ecbd*, 60 nM NDH2, 50 mM menaquinone-1 in 50 mM MES pH 6.5, 150 mM NaCl, and 3 µM of the inhibitor hits. Hit confirmation was performed by measuring the NADH oxidation rate on a FLUOstar system (250 µM NADH). The data was analyzed by ANOVA with Dunetts *post hoc* test (Graphpad Prism). Significance was set at p<0.05. This yielded compound **ENL1148** (Z57044282 Enamine) as the most promising hit. The IC_50_ value and inhibitor competition was determined using the *Mtbd* or *Ecbd* oxygen consumption rate on an Oxygraph system (Hansatech) after addition of 500 µM NADH.

### NDH2 activity counter-screen

Hits were counter-screened against NDH2 to ensure the observed inhibitory effect is caused by direct inhibition of *Mtbd*. Briefly, NADH oxidation was monitored in a solution of 100 nM NDH2 with 200 µM menadione in 50 mM MES pH 6.5, 150 mM NaCl, 0.005% LMNG. NDH2 turnover was initiated by addition of 150 µM NADH, followed by rate determination. The data was analyzed by ANOVA with Dunetts *post hoc* test (Graphpad Prism). Significance was set at p<0.05.

### Growth inhibition in *M. tuberculosis* H37ra

Inhibitor growth inhibition was determined using the *M. tuberculosis* H37ra strain. Cells were grown in 7H9 media supplemented with ADS (0.5% BSA, 0.75% dextrose, and 0.08% NaCl). The cells were diluted to OD 0.1 prior to exposure to the desired inhibitor concentration. H37ra cells were exposed to **Q203** or **WSL017**. To determine synergistic effects between cyt *bcc:aa*_*3*_ and the cyt *bd* inhibitors, cells were exposed to a combination of **Q203** and **WSL017**. Sterile media and DMSO controls were included to ensure assay validity. Cells were incubated for 7 days in a humidified incubator at 37ºC. Growth was quantified by addition of Alamar blue dye (Thermo Fisher) (5 vol% final) followed by incubation for 6-8 hours at 37ºC. Alamar blue fluorescence was determined by fluorescence (530 ex, 590 em.) and normalized to the DMSO control to measure cell growth to calculate the percentage of cell growth.

### Cryo-EM Sample Preparation and Data Collection

UltrAUfoil R1.2/1.3-Cu 300 mesh grids were freshly glow discharged at 15 mA for 45 s followed by 15 mA and 90 S on the foil side. LMNG solubilized *Ecbd* was incubated with 100 µM **WSL017**, followed by the application of 4 µl sample (1.5 mg/mL) on the grid. Sample blotting was performed for 6 seconds, at 20 blot force using a vitrobot IV device (Thermo Fischer) operating at 4 °C and 100% humidity, directly before plunge freezing in liquid ethane.

A total of 8645 movies were collected on a Titan Glacios (Thermo Fisher) operating at 200 keV, equipped with a Falcon 4i detector and selectris energy filter with a slit width of 20 eV. Movies were acquired using a total dose of 100 *e*/Å^2^ with 100 frames, at 130,000× magnification with a calibrated pixel size of 0.880 Å and a defocus range of -1.0 to -2.0 µm.

### Cryo-EM data analysis

Data processing was carried out in CryoSPARC. Image preprocessing was performed in CryoSPARC Live^42^ using Patch Motion Correction and Patch CTF estimation. Initial particle picking was conducted with blob picking, followed by template generation for both monomeric and dimeric particles. The processed micrographs and templates were then exported to CryoSPARC for downstream analysis. Template-based particle picking was applied to both monomer and dimer datasets, followed by multiple rounds of 2D classification to remove junk particles. Three-volume unsupervised *ab initio* reconstruction was performed to generate initial and bait volumes for further particle cleanup. Successive rounds of heterogeneous refinement were carried out until a consistent class containing clean particles was obtained. This refined particle stack was subjected to Non-uniform refinement^43^, including local CTF refinement and higher-order aberration fitting. To mitigate beam damage, movies were reprocessed using patch-based motion correction with the final 50 frames excluded. Particles were then subjected to reference-based motion correction^44^, followed by an additional round of non-uniform refinement to achieve the final reconstructions.

## Data availability

The cryo-EM map is deposited at the Electron Microscopy Data Bank under accession code: EMD-55645. Atomic model of *Ecbd* bound to **WSL017** has been deposited to the Protein Data Bank under accession number: 9T7N. All other data are presented in the main text or supplementary information.

## Acknowledgements

We would like to thank Dr. Dirk Bald (VU Amsterdam, The Netherlands) for providing the MB43 cells and the pET17b_CydABX_Linkerstreptag construct for *Ecbd* expression. Furthermore, we would like to thank Prof. Dr. Gregory Cook (University of Otago, New Zealand) for providing the MC^2^ 155 ΔCydAB strain and pLHCyd plasmid for *Mtbd* expression. S.M.H. and A.H. acknowledge the Dutch Research Council (NWO) for funding through a VIDI grant (VI.Vidi.213.057).

## Author contributions

Conceptualization: LJCJ, TTVDV, SMH

Methodology: TTVDV, AH, JPVP, SMH, LJCJ

Investigation: TTVDV, AH, WSL

Visualization: TTVDV

Funding acquisition: LJCJ

Project administration: LJCJ

Supervision: LJCJ, SMH

Writing – original draft: TTVDV, LJCJ

Writing – review & editing: TTVDV, AH, JPVP, SMH, LJCJ

Data curation: TTVDV

## Conflict of interest

The authors declare that they have no conflicts of interest.

## Abbreviations

AurDyp: Aurachin D, Cyt *bd*, Cytochrome *bd*
Ecbd: E. coli cytochrome *bd*-I
LMNG: Lauryl Maltose Neopentyl Glycol
MK: Menaquinone
*Mtbd*: *M. tuberculosis* cytochrome *bd*
NDH2: NADH dehydrogenase type-2
UQ: Ubiquinone-10; TMS, tetramethylsilane
TLC: Thin layer chromatography
LC-MS: Liquid Chromatography-Mass Spectrometry
NMR: Nuclear Magnetic Resonance

## Notes

### Competing Interest Statement

The authors have declared no competing interest.

