## Supplementary Information for "Discovery of a novel chemotype targeting *Mycobacterium tuberculosis* cytochrome *bd* through rapid screening and structural elucidation"

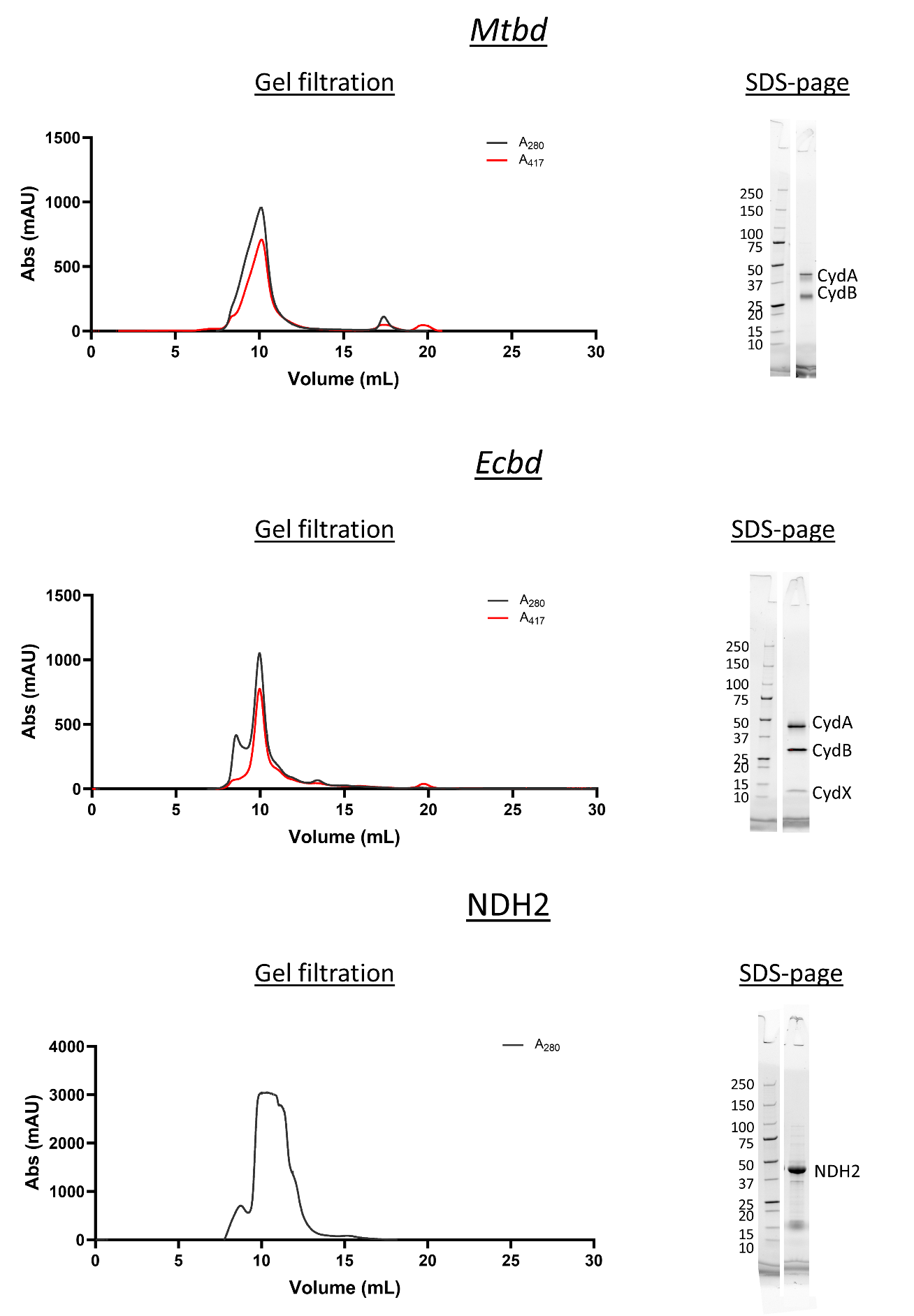

**Figure S1**. Gel filtration and SDS-Page (TGX stain) of the purified Mtbd, Ecbd and NDH2.

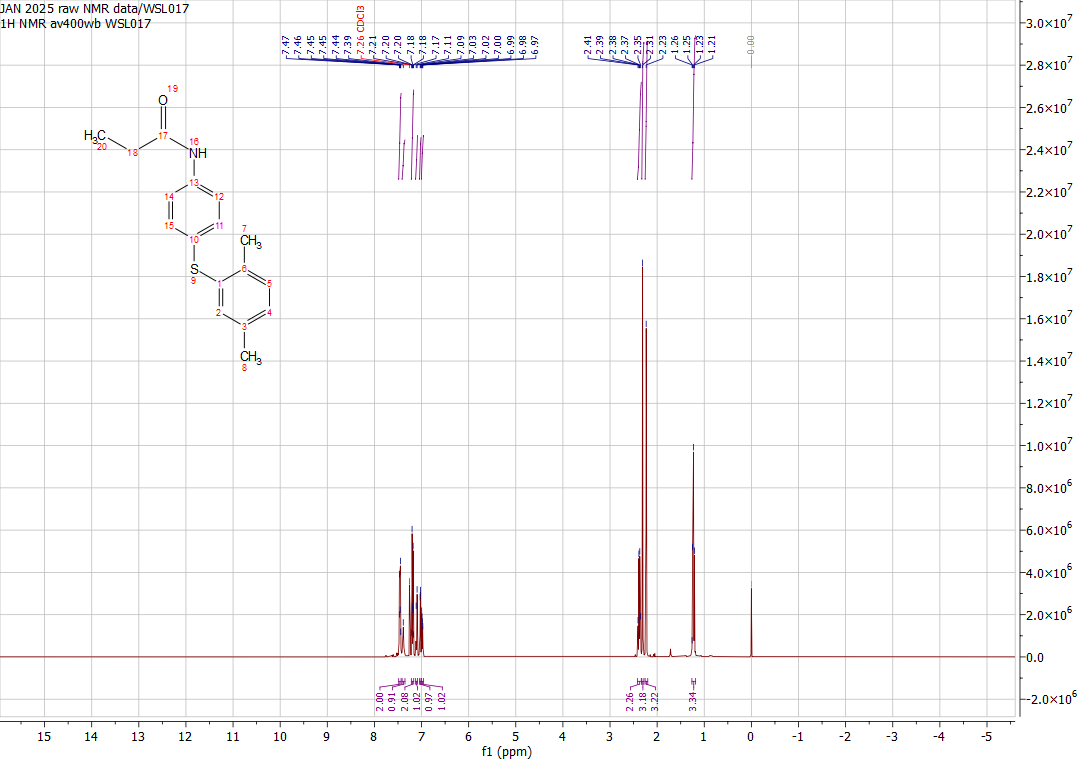

**Figure S2.** ^1^H NMR of N-(4-((2,5-dimethylphenyl)thio)phenyl)propionamide (**WSL017**)

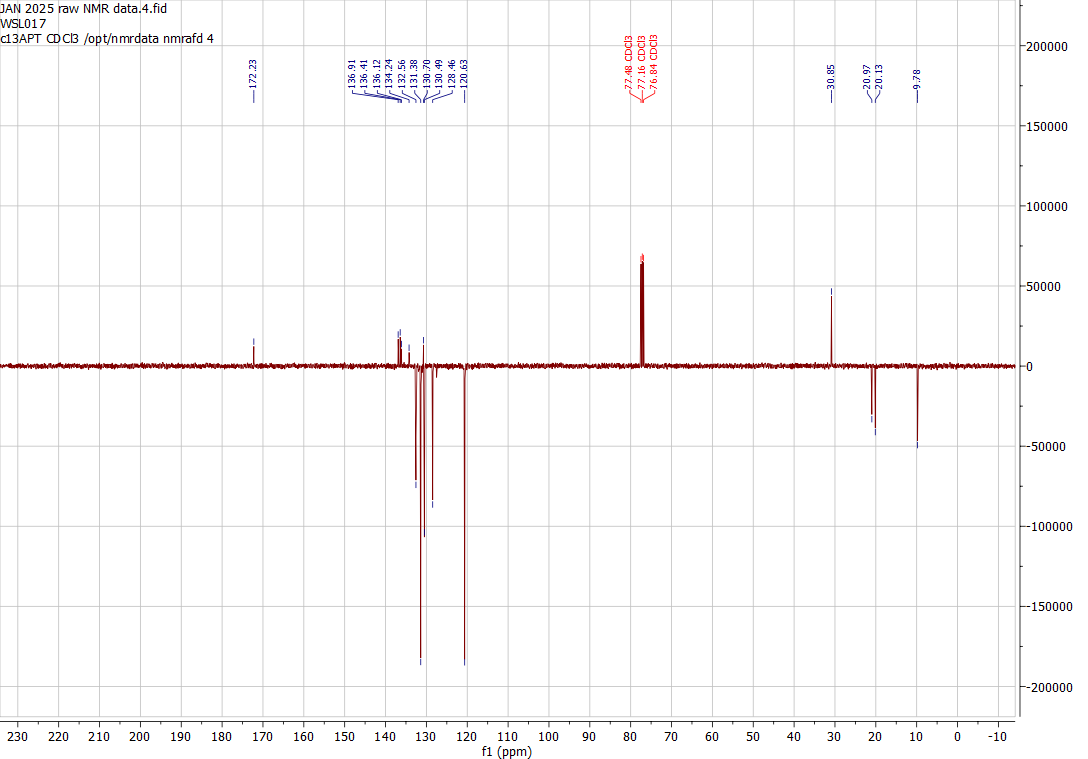

**Figure S3.** ^13^C NMR of N-(4-((2,5-dimethylphenyl)thio)phenyl)propionamide (**WSL017**)

**Table 1**. Screening results and hit confirmation in detergent and proteoliposomes

| **Compound number (ENL library)** | **SMILES** | **Z-score** | **Hit confirmation detergent**  **(P value)** | **Hit confirmation Liposomes (P value)** |
| --- | --- | --- | --- | --- |
| 496 | CC(NC(=O)C=C)C1(CC1)S(=O)(=O)C | 4.42 | 0.0001 | 0.9991 |
| 573 | C=CC(=O)N1CCC(CC1)C(=O)N2CCC(CC=3N=CON3)CC2 | 3.23 | 0.0008 | 0.9998 |
| 575 | CCN(CC(=O)NC1CCN(C(=O)C)C=2C=CC=CC12)C(=O)C=C | 4.47 | 0.5493 | 0.94 |
| 576 | OC1CCCN(C1)C(=O)C2(CCN(CC2)C(=O)C=C)C=3C=CC=CC3 | 4.62 | 0.0114 | 0.9999 |
| 653 | CCN(CC=1C=CC=C2OCOC21)C(=O)C=C | 4.22 | 0.0009 | 0.9999 |
| 778 | OCCNC(=O)C1CCN(CC1)C(=O)C=C | 3.36 | 0.0001 | 0.2583 |
| 964 | CC=1C=CC(SC=2C=CC(NC(=O)CCl)=CC2)=CC1C | 3.85 | 0.0001 | 0.2583 |
| 1148 | CC=1C=CC(C)=C(SC=2C=CC(NC(=O)CCl)=CC2)C1 | 3.69 | 0.0001 | 0.1612 |
| 1377 | FC(F)(F)CC(=O)N1CCN(CC1)C(=O)CCl | 3.01 | 0.0014 | 0.9991 |
| 1413 | ClCC(=O)NC1CN(C1)S(=O)(=O)C=2C=CC=3OCCOC3C2 | 3.18 | 0.0006 | 0.9993 |
| 1513 | CC(Cl)C(=O)N1CC(C)CN(CC=2C=CC=CC2)CC1C | 3.56 | 0.015 | 0.999 |
| 1515 | CC(Cl)C(=O)N1CC(C1)N2C=C(N=N2)C(=O)N | 4.53 | 0.0006 | 0.9994 |
| 1516 | CC(Cl)C(=O)N(CC1=CN=CS1)C2CCN(CC2)C(=O)C | 3.59 | 0.0011 | 0.9986 |
| 1594 | CC(Cl)C(=O)NC=1C=CC(C)=C(C1)S(=O)(=O)N2CCOCC2 | 3.11 | 0.0031 | 0.9986 |
| 1595 | ClCC(=O)N(CC1=CC=CO1)CC=2C=CC=CC2 | 3.43 | 0.0003 | 0.9999 |
| 1596 | CC(Cl)C(=O)NC=1C=CC=C(Cl)C1 | 4.55 | 0.002 | 0.9994 |
| 2165 | CC=1C=CC=CC1C(NC(=O)CCl)C=2C=CC=CC2C | 9.09 | 0.0001 | 0.9996 |
| 2570 | COC=1C=CC(C=C(C#N)C(=O)NC=2C=CC(=CC2)[N+](=O)[O-])=CC1 | 11.72 | 0.0001 | 0.9863 |
| 3053 | CC=1C=CC(C=C(C#N)C(=O)NC=2C=CC=C3N=CC=CC23)=CC1 | 3.80 | 0.1058 | 0.9812 |
| 3075 | CC1=NN(C)C(C)=C1C=C(C#N)C(=O)NC=2C=CC=CC2Cl | 3.23 | 0.0048 | 0.9998 |
| 3076 | CCCCNC(=O)C(=CC=1C=CC=C(Cl)C1Cl)C#N | 3.01 | 0.0036 | 0.9998 |
| 3084 | CCCCNC(=O)C(=CC=1C=CC(Cl)=CC1)C#N | 3.03 | 0.0114 | 0.9993 |
| 3153 | NC(=O)C(=CC1=CSC(COC=2C=CC=CC2)=N1)C#N | 3.10 | 0.0014 | 0.998 |
| 3155 | CC(=CC=1C=CC=CC1)C=C(C#N)C(=O)NC2CC2 | 3.16 | 0.0329 | 0.998 |
| 3156 | CC1=NC=CN1CC=2C=CC(NC(=O)C(=CC3=CC=CO3)C#N)=CC2 | 3.45 | 0.0012 | 0.9991 |
| 3207 | ClCC(=O)NCC=1C=CC(=CC1)C(=O)N2CCOCC2 | 3.24 | 0.0003 | 0.9863 |
| 3374 | CS(=O)(=O)C=1C=CC(CNC(=O)CCl)=CC1 | 7.57 | 0.0099 | 0.6406 |
| 3501 | ClCC(=O)N1CCN(CC1)S(=O)(=O)C=2C=CC=3OCCOC3C2 | 7.07 | 0.0031 | 0.9995 |
| 3726 | CN1N=CC=C1CN2C[C@@H](F)C[C@H]2CNC(=O)CCl | 4.11 | 0.0048 | 0.9993 |
| 3929 | CN1N=CC=C1C(CO)NC(=O)CCl | 3.24 | 0.0017 | 0.9988 |
| 4797 | CC1=NC(=NO1)C=2C=CC(CNC(=O)C=C)=CC2 | 3.92 | 0.0006 | 0.9812 |
| 4997 | CC=1C=CC=C(C1)C2(CCC2)NC(=O)C=C | 5.65 | 0.0011 | 0.9988 |
| 5038 | CN1C=C(N=N1)C(C)(C)NC(=O)C=C | 3.20 | 0.0036 | 0.9994 |
| 5085 | C=CC(=O)NC1C2CCOC2C31CCCC3 | 3.55 | 0.0131 | 0.9997 |
| 5676 | C=CC(=O)NC=1N=C2N=CC=CN2N1 | 4.40 | 0.0224 | 0.9986 |
| 6185 | CC1(CCCN1C(=O)C=C)C=2C=CC=CC2 | 5.97 | 0.0086 | 0.9988 |
| 6717 | C=CC(=O)NC1CCN2CCCC2C1 | 4.23 | 0.0196 | 0.6696 |
| 6820 | FS(=O)(=O)N1CCN(CC1)C=2C=CC=CC2 | 4.34 | 0.0476 | 0.6406 |
| 7026 | CC(=O)N1CCC2(CC1)CCC=3C=CC(OS(=O)(=O)F)=CC3O2 | 3.01 | 0.0255 | 0.0487 |
| 7182 | CC=1C=C(C(=O)NC=2C=CC(OS(=O)(=O)F)=CC2)C(C)=NC1C | 8.49 | 0.0114 | 0.1483 |
| 7502 | OC(=O)C=1C=CC=CC1NC(=O)C(=CC=2C=CSC2)C#N | 3.61 | 0.0002 | 0.4968 |
| 7527 | FC=1C=CC(CNC(=O)C(=CC=2C=CC=CC2)C#N)=CC1 | 3.51 | 0.0001 | 0.0721 |
| 7614 | FC=1C=CC(CCNC(=O)C(=CC2=CC=CO2)C#N)=C(F)C1 | 3.24 | 0.0086 | 0.998 |
| 7642 | ClC=1C=CC(=CC1)N2C=C(C=C(C#N)C(=O)NCC=C)C=N2 | 3.22 | 0.0001 | 0.0358 |
| 7725 | O=C(NC1CC1)C(=CC=2C=CC(=CC2)N3CCCCC3)C#N | 3.72 | 0.0099 | 0.9984 |
| 8695 | CC#CC(=O)N(C)CC1=NC(=CS1)C(F)(F)F | 13.52 | 0.0114 | 0.9982 |
| 9524 | CC(Cl)C(=O)N1CC2(CCOCC2)C1C3CC3 | 6.99 | 0.0373 | 0.8525 |
| 9953 | OB(O)C=1C=C2C=CC=CC2=NC1O | 3.24 | 0.0086 | 0.9996 |
| 10255 | CCS(=O)(=O)N1CCC(CB2OC(C)(C)C(C)(C)O2)C1 | 3.64 | 0.0027 | 0.9997 |
| 10257 | CC1(C)OB(OC1(C)C)C2=CCN(CC2)C(=O)C=3C=CC=C4C=CN=CC34 | 3.87 | 0.0224 | 0.9996 |
| 10333 | CC(C)NC(=O)NC=1C=CC=CC1NC(=O)C(F)Cl | 3.77 | 0.2564 | 0.9995 |
| 10334 | CC(NC(=O)C(F)Cl)C1=NC(=CS1)C=2C=CC=CC2 | 3.16 | 0.0604 | 0.9993 |

**Table 2**. Cryo-EM data collection, refinement and validation statistics.

|  | ***Ecbd* dimer WSL017 (EMDB-55645, PDB 9T7N)** |
| --- | --- |
| **Data collection and processing** |  |
| Magnification | 130,000 |
| Voltage (kV) | 200 |
| Electron exposure (e–/Å^2^) | 100 |
| Defocus range (μm) | -0.8 - -2.0 |
| Pixel size (Å) | 0.880 |
| Symmetry imposed | C2 |
| Initial particle images (no.) | 5,721,267 |
| Final particle images (no.) | 193,848 |
| Map resolution (Å) | 2.60 |
| FSC threshold | 0.143 |
| Map resolution range (Å) | 2.5-2.7 |
| **Refinement** |  |
| Initial model used (PDB code) | 9RZV |
| Model resolution (Å) | 2.6 |
| FSC threshold | 0.143 |
| Map sharpening *B* factor (Å^2^) | -60 |
| **Model composition** |  |
| Non-hydrogen atoms | 15849 |
| Protein residues | 1902 |
| Ligands | 4 HEM, 2 A1JN4, 6 LPP, 2 PGT, 4 UQ8, 2 OXY, 2 A1JT9 |
| Water | 15 |
| ***B* factors (Å^2^)** |  |
| Protein | 112.53 |
| Ligand | 117.20 |
| **R.m.s. deviations** |  |
| Bond lengths (Å) | 0.009 |
| Bond angles (°) | 0.83 |
| **Validation** |  |
| MolProbity score | 1.24 |
| Clashscore | 4.69 |
| Poor rotamers (%) | 0.39 |
| **Ramachandran plot** |  |
| Favored (%) | 98.36 |
| Allowed (%) | 1.64 |
| Disallowed (%) | 0.00 |

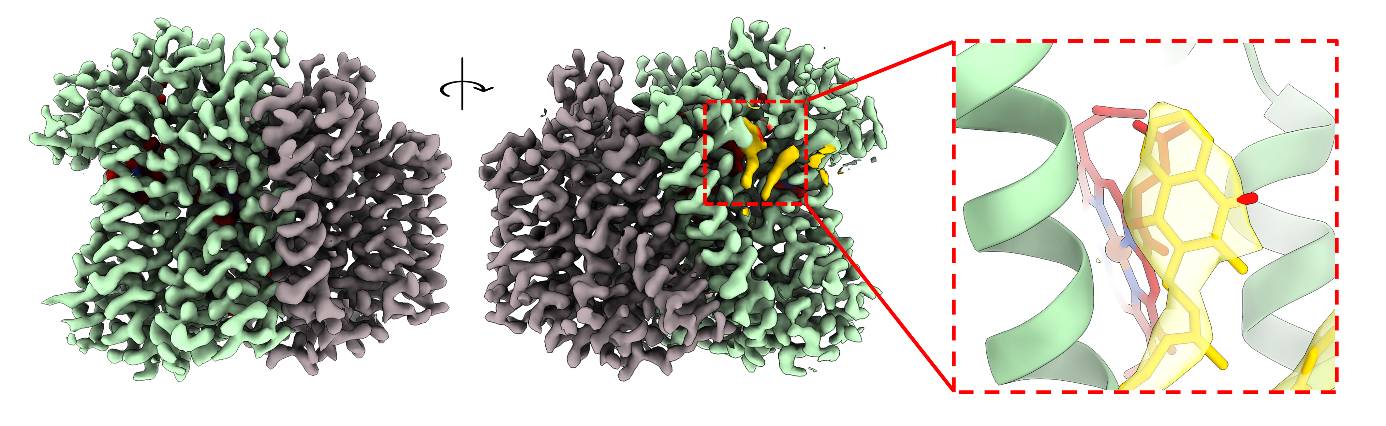

**Figure S4**. Colored cryo-EM map of Mtbd resolved in the presence of **WSL017** (3.1 Å). CydA is shown in green with CydB in grey. No change in the density of MK-9 is observed.

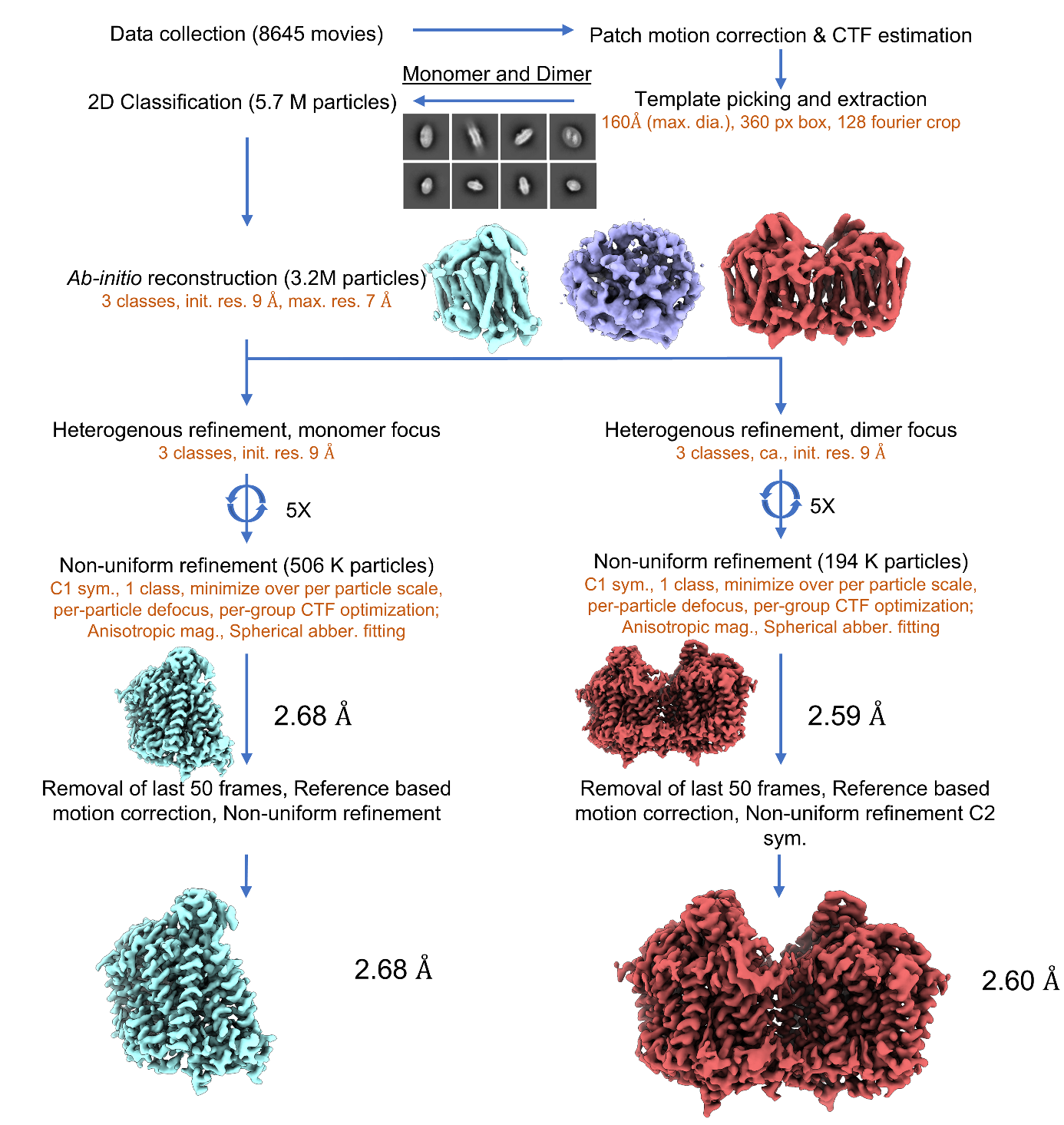

**Figure S5**. Cryo-EM data processing procedure of **WSL017** bound to Ecbd.

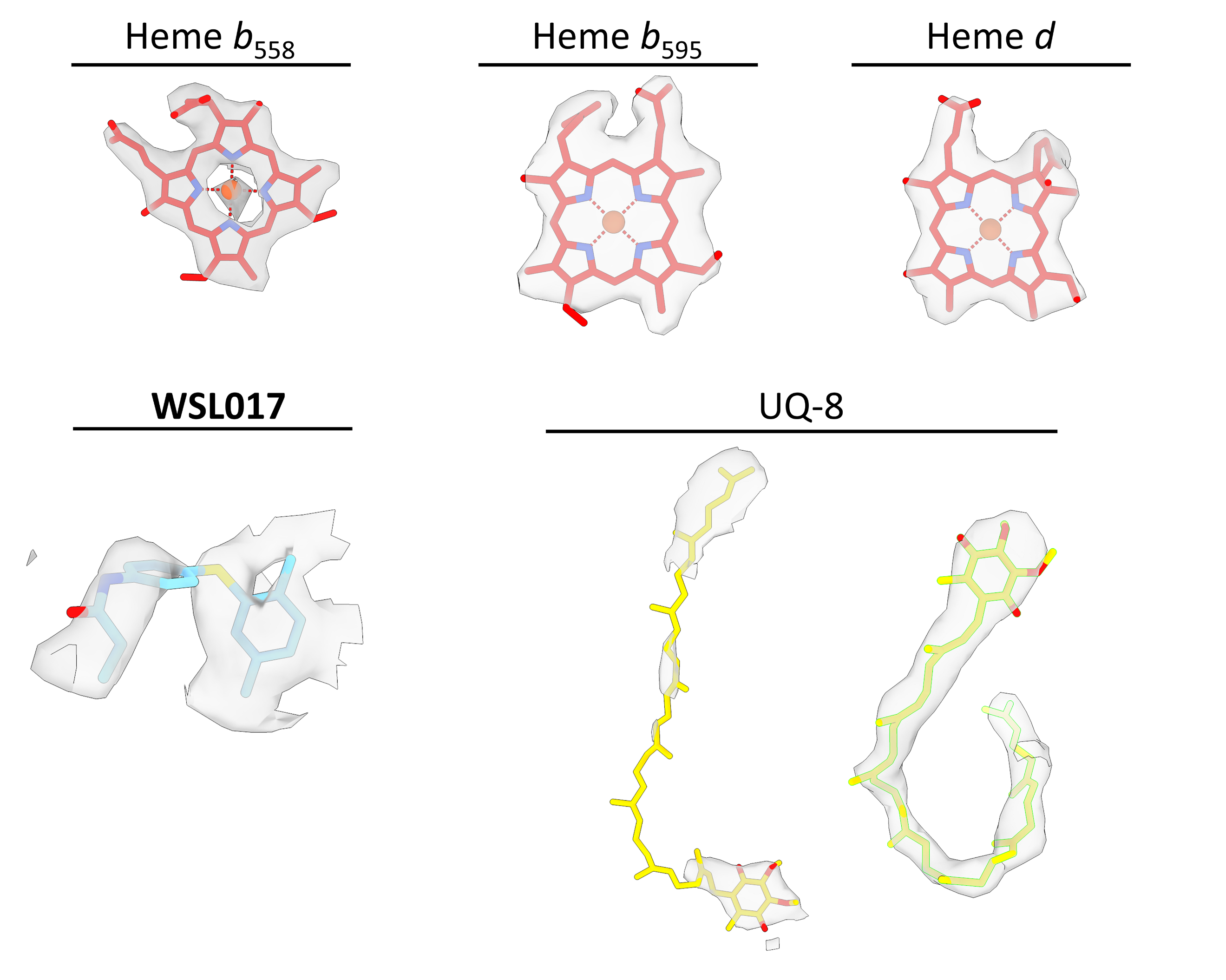

**Figure S6**. Density features of the main ligands bound to Ecbd.

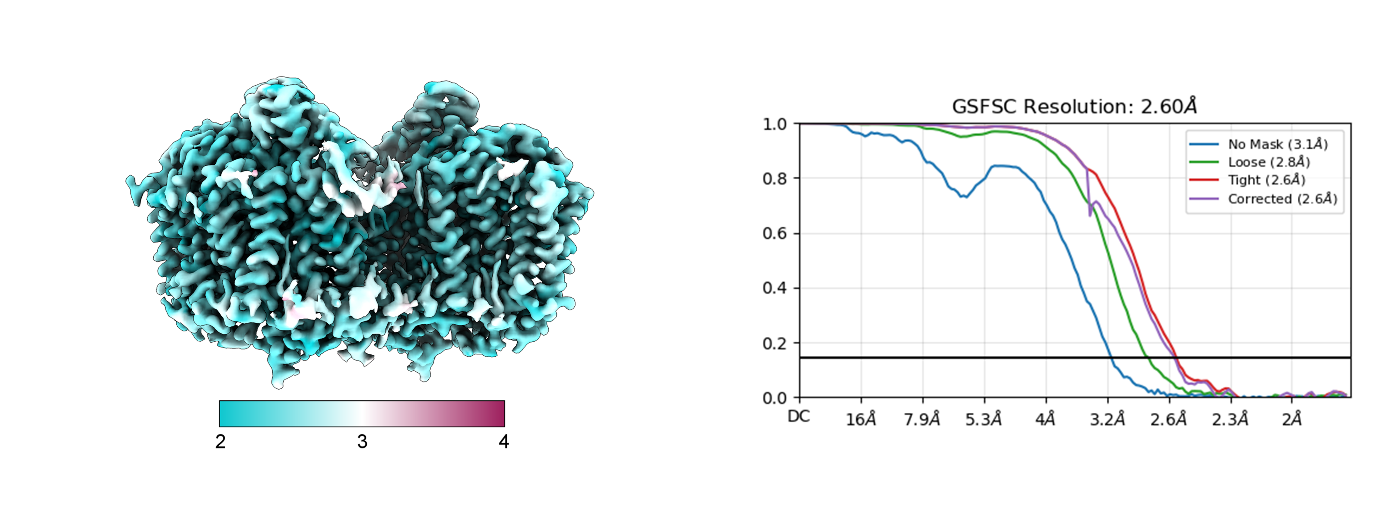

**Figure S7**. Local resolution and fsc curve of the Ecbd dimer structure bound to **WSL017**.

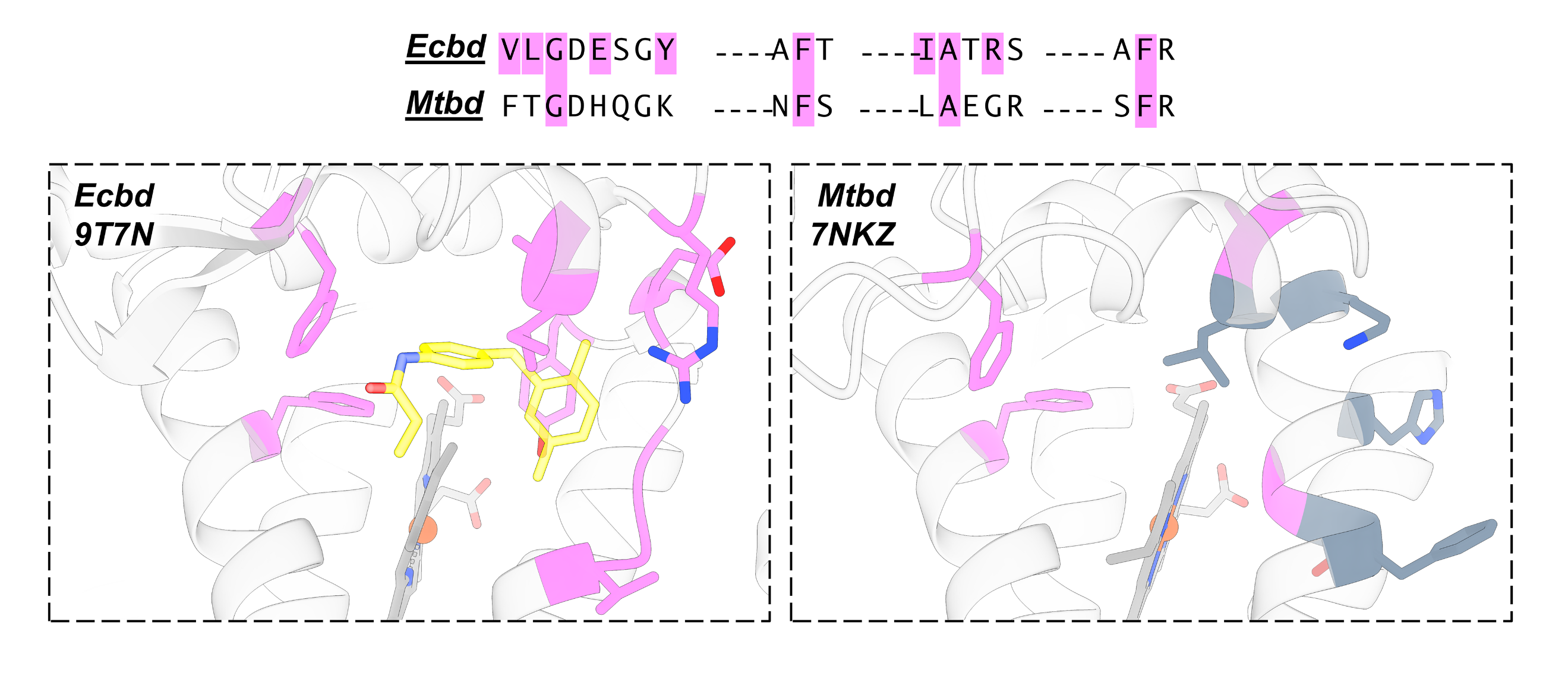

**Figure S8**. Comparison of the **WSL017** binding pocket in Ecbd, highlighting the interacting residues in pink, with the equivalent region in Mtbd. The corresponding residues in Mtbd are shown with conserved interactions highlighted in pink.

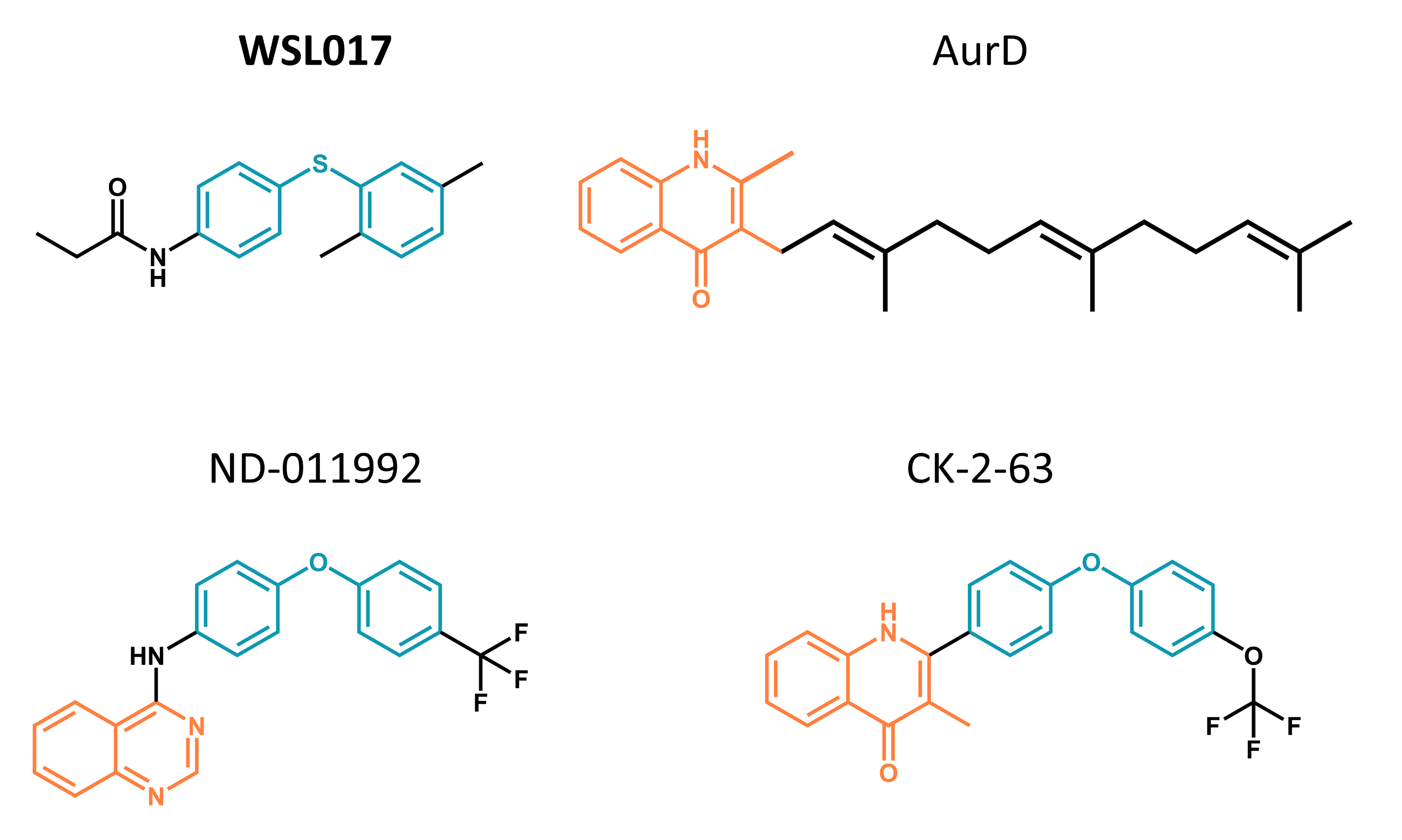

**Figure S9**. Overview of common cytochrome bd inhibitor features. The aurachin D (AurD) headgroup is highlighted in orange, while the **WSL017** backbone is highlighted in blue.

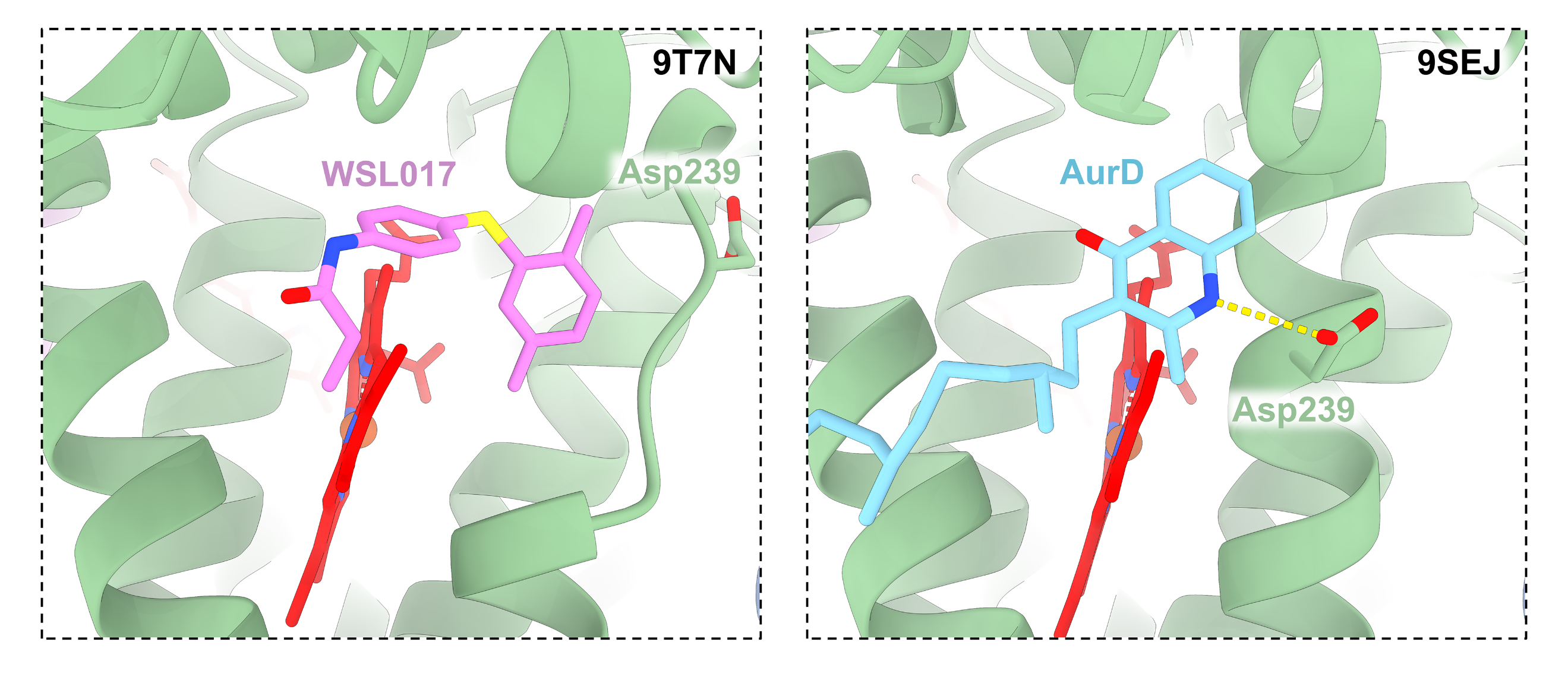

**Figure S10**. Different binding mode of **WSL017** and AurD in the Ecbd quinol oxidation site. AurD induced active site refolding by hydrogen bonding with Asp239^CydA^.
